# Novel origin of replication for environmentally isolated *Pantoea* strain enables expression of heterologous proteins, pathways and products

**DOI:** 10.1101/2025.05.10.653210

**Authors:** Alex Codik, Ankita Kothari, Hualan Liu, Benjamin L. Weinberg, Trenton K. Owens, Aparajitha Srinvasan, Alex Rivier, Thomas Eng, Adam P. Arkin, Adam M. Deutschbauer, Aindrila Mukhopadhyay

**Affiliations:** Biological Systems and Engineering Division, Lawrence Berkeley National Laboratory, Berkeley, CA 94720, USA; Comparative Biochemistry Graduate Group, University of California, Berkeley, Berkeley, CA 94720, USA; US Department of Energy Joint Genome Institute, Lawrence Berkeley National Laboratory, Berkeley, CA 94720, USA; Environmental Genomics and Systems Biology Division, Lawrence Berkeley National Laboratory, Berkeley, CA 94720, USA; Department of Plant and Microbial Biology, University of California, Berkeley, CA 94720, USA; Department of Bioengineering, University of California, Berkeley, CA 94720, USA

**Keywords:** *Pantoea*, origin of replication, magic pool, non-model bacteria, isoprenol, indigoidine, bioproduction

## Abstract

Plasmids isolated or characterized from environmental samples serve as a resource that can be used to develop genetic tools for characterizing recently isolated or less-studied microbes. In this report, we leveraged sequences from a previously characterized groundwater plasmidome and developed a screen to identify novel plasmid origins. Putative origin sequences were used to construct a barcoded plasmid library, which contained both known and newly predicted origins. This library was tested against a panel of representative bacterial strains and led to the identification of 3 novel origins that putatively replicate in gram-negative bacteria not previously associated with these origin sequences. We empirically validated one of the newly identified origins, 6911, to be functional in both the model bacterial strain, *Escherichia coli* BW25113, as well as in *Pantoea* sp. MT58, a fast growing and metal tolerant, environmentally important bacterium from the widespread *Pantoea* genus. We confirmed that a plasmid bearing origin 6911 as the sole origin could replicate and had a copy number of 9 (± 2) in *Pantoea* sp. MT58. We successfully used a plasmid based on the new origin to express the reporter protein GFP, and two non-native metabolite pathways for the natural product, indigoidine and the terpenoid compound, isoprenol. By pairing functional novel origins of replication to non-model organisms this pipeline can expand the tool kit for genetic manipulations of both model and less-studied bacteria.

Abstract figure.We developed a host-origin pair identification pipeline by constructing a plasmid library containing both literature-sourced and computationally predicted origins of replication. By conjugating this library into non-model microbes, we identified functional origins in diverse bacteria, and could leverage these findings to develop genetically tractable hosts for microbial engineering.

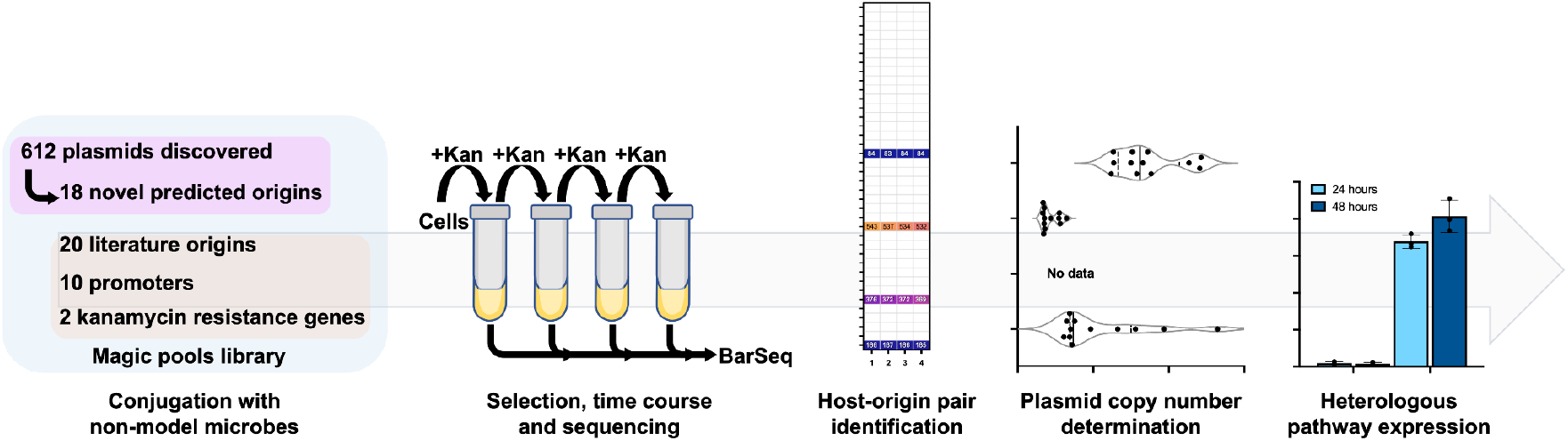

## 1. Introduction

Genetic transformation is a critical step to transition from a genomics-based examination of a bacterial strain to genetics. Transformation of a plasmid capable of replicating in a bacterial host enables an array of genetic approaches that are indispensable in characterizing or engineering microbial strains. However, it can be a challenge to find and/or construct a plasmid that is able to self-replicate in a microbe of interest [1–5]. Specifically, the plasmid’s origin of replication is a key requirement but predicting origins that will function in a given host is challenging [6]. However, discovery of new origins that function in diverse, non-model microbes has not been extensively explored.

Recent advances have allowed for the identification of many features of plasmids that are valuable in understanding the functional role of a plasmid as a mobile genetic element in the environment [7–12], and to reveal biological parts that can be used in molecular biology approaches [11–15]. However, identification of the sequence that encodes the origin of replication is challenging and design of origins *de novo* remains elusive. One path forward is the use of existing plasmid sequences that can be analyzed to predict putative origin sequences and can then be experimentally verified. A plasmidome is the total collection of bulk plasmids present in a given environment that can be measured by whole genome sequencing (WGS) [16]. Targeted plasmidome studies, as well as metagenomics studies, reveal a vast range of native plasmids that provide a deep resource to address this challenge [17–20]. In this report, we leveraged a previously characterised plasmidome from groundwater samples from the Oak Ridge Field Research Center (ORFRC) containing over 600 unique plasmid sequences [17]. The best available tool for predicting origins is DoriC [8]. Though primarily targeted at replicating sequences in the genome, this workflow can also be applied to plasmid sequences. A subset of these groundwater plasmidome sequences were analyzed using the DoriC software and 18 putative novel origins of replication were predicted [8]. We then adopted a magic pool approach to build a DNA barcoded plasmid library containing the predicted origin sequences [21].

A primary goal of our study was to discover origins that would enable identification of new host-plasmid pairings and advance the usable genetic tools in new environmentally and biotechnologically important bacterial strains. As such we screened the plasmid pool containing a combination of putative and known origins against a well-studied model bacteria (e.g. *E. coli* BW25113) and less-studied environmental strains (e.g. *Pantoea* sp. MT58). The *Pantoea* genus is especially important for further studies; it is a widely distributed genus found both in plant and soil associated communities spanning pathogenic [22–25] and commensal [23,26] species. *Pantoea* sp. are also seen as a valuable new platform for biotechnology applications [14,23,27–29]. In both of these contexts deeper genetic tools and an available model-system, such as the *Pantoea* sp. MT58 could be valuable. In the ORFRC, *Pantoea* sp. MT58 is member of the sediment microbial community that is gaining interest due to its metal tolerance capabilities [30], and a recent pangenome study for this bacterium highlighted its facultative anaerobic capabilities being involved in survival in the contaminated groundwater at the ORFRC [31]. In our study we sought to discover additional genetic parts for use in this emerging microbe of interest and demonstrate the use of new plasmids for expression of proteins and pathways. Our discovery workflow and extensive characterization of a new origin of replication for *Pantoea* sp. MT58 is described in the following sections.

## 2. Materials and Methods

### 2.1 General methods, bacterial cultures, and growth media

The bacterial strains and plasmids used in this study are listed in Data S1. *E. coli* BW25113 was acquired from the Joint BioEnergy Institute’s registry (https://public-registry.jbei.org/) [32]. All other strains listed in Table 1 were isolated from the Oak Ridge Field Research Center (ORFRC); (https://public.ornl.gov/orifc/orfrc3_site.cfm).

Competent cells of *E. coli* TransforMax EC100D *pir*-116 cloning strain and the *E. coli* WM3064 conjugation donor strain were purchased from Lucigen (Biosearch Technologies, Alexandria, MN). *E. coli* 10-β competent cells were purchased from New England BioLabs (NEB, Ipswich, MA, USA). All bacterial strains were grown in liquid and solid medium using a Luria-Bertani (LB) base (Becton Dickinson, Milpitas, CA). Antibiotic concentrations for selection were carbenicillin (carb, 50 µg/mL) and kanamycin (kan, 50 µg/mL). When the *E. coli* WM3064 strain was used, media was supplemented with diaminopimelic acid (DAP) at a concentration of 300 µM.

All non-magic pool plasmids were constructed using standard Gibson cloning methods [33]. Plasmid DNA isolation was performed using a miniprep plasmid isolation kit according to the manufacturer’s instructions (Qiagen, Redwood City, CA). General microbiological manipulations, PCR amplification, golden gate assembly, cloning, and transformation procedures were as described previously [21]. Enzymes were purchased from New England Biolabs (NEB, Ipswich, MA, USA) and Thermo Fisher Scientific (Waltham, MA, USA). Oligonucleotide primers were obtained from Integrated DNA Technologies (IDT, Coralville, IA, USA). Double strand DNA gene fragments were ordered for synthesis from Genewiz Inc (Azenta Life Sciences, South Planefield, NJ). All plasmids constructed, not including magic pool library construction, were verified by whole plasmid sequencing by Plasmidsaurus Inc (South San Francisco, CA). Unless noted, all reagents and media components were purchased from Sigma-Aldrich (St. Louis, MO, USA).

### 2.2 Origin of replication prediction

Of the roughly 600 circular plasmids from our earlier plasmidome study, 32 circular plasmid sequences were shared with DoriC for curation of plasmid-based origin predicted [8]. Of the 32 plasmids that were shared, 18 predicted origins of replication were provided by DoriC [8]. The resulting sequences were used in the resulting work.

### 2.3 Construction of the kanamycin magic pool origin libraries

Part 1 origin plasmids were synthesized into a pJW52 backbone [33] by Genscript USA Inc (Piscataway, NJ). Parts 2 (promoter for drug resistance marker) and 3 (drug resistance coding sequence (CDS)) (also in pJW52 backbones) plasmids were constructed previously [21]. All plasmids containing parts 1, 2, and 3 can be found in Data S2. For all part 1 origins, 100 ng of each plasmid were added to a part 1 mix. This was also done for parts 2 and 3, separately. 100 ng of each mix was then added into the golden gate assembly [34]. The origin magic pool was constructed by golden gate assembly using methods described previously [21].

The magic pool was built in two dilutions, “1x” and “2x” based on the number of *E. coli pir*+ colonies added to 100 mL of LB medium supplemented with DAP and carbenicillin. The magic pool was allowed to grow overnight, followed by a glycerol stock and four plasmid DNA isolations per dilution. An amplification of the barcoded region of each plasmid assembly was performed to check the number of barcodes per assembly. For the magic pool the “1x” dilution was selected due to its barcode coverage relative to the total combination of possible vectors (∼10x). Utilizing the DNA isolated previously, the magic pool was transformed into the *E. coli* WM3064 conjugation donor strain. The entirety of the transformation was then grown in 100 mL of media supplemented with DAP and stored via glycerol stock after an overnight incubation.

The plasmid DNA that was isolated from the *E. coli pir*+ cells, and subsequently used to transform into the *E. coli* WM3064 conjugation donor, was sequenced by Plasmidsaurus using long-read sequencing (once with PacBio sequencing technology and once with in-house using Nanopore sequencing) with custom analysis and annotation. Sequencing data was analyzed by searching through each long read and comparing the sequence to a list of parts that were used in the construction. The output of this search was a list of barcodes and the parts (1, 2 and 3) associated with the barcode. For us to confidently assign a unique barcode to an origin part, we had to observe at least two unique long-reads that supported this observation. In some cases a single barcode was associated with multiple parts from each grouping of parts (i.e. multiple part 1s can be associated with a single barcode) (see Figure S1 and Data S3 for part 1 barcode-origin associations). These non-unique barcodes were not used for any further analysis.

Part 2.4 showed a 160 bp sequence similarity to the part 4 backbone (promoter driving *bla* gene). This caused the software that analyzed the long read sequencing data to not be able to differentiate between part 2.4 and the backbone. In Figures S2-S4 “unmapped part 2” represents either unmapped part 2s or part 2.4.

### 2.4 Magic pool conjugation

The following protocol was adapted from [21]. Recipient strains were streaked out on LB agar for isolated colonies. A single colony of the recipient was used to inoculate a 10 mL volume of LB and grown overnight at 30 °C and shaking at 200 rpm. A 2 mL glycerol stock of the *E. coli* WM3064 donor strain, containing the magic pool library, was inoculated into 50 mL of LB supplemented with carbenicillin and DAP and allowed to grow at 37 °C, while shaking at 200 rpm, for 4 hours. 1 OD of donor cells were pelleted in a microcentrifuge tube and washed twice with LB. 1 OD of recipient cells were pelleted and washed once with LB. Each conjugation contained a 1:1 OD ratio of donor:recipient cells. A donor pellet and a recipient pellet were resuspended in the same 200 µL of LB and plated onto an LB plate supplemented with DAP. Conjugations were incubated at 30 °C overnight. Cells were harvested from the conjugation plates and resuspended in 4 mL of LB with 15% glycerol. These conjugation stocks were frozen at −80 °C.

To determine the number of colonies per mL, a tube of conjugation stock was thawed and serially diluted down to 1×10^−6^. Dilutions 1×10^0^ to 1×10^−4^ were plated on kanamycin selective media while dilutions 1×10^−5^ and 1×10^−6^ were plated on non-selective media to check for transformation efficiency and for survival of the recipient. Colony counts from these plates were used to estimate the volume of conjugation mix would be needed to get ∼5,000 colonies. This volume was then either inoculated directly into liquid selection, and grown for the time course experiments, or plated onto LB supplemented with kanamycin and incubated overnight at 30 °C to form small colonies.

For the liquid selection time course experiment, ∼5,000 CFUs were inoculated into 10 mL of LB supplemented with kanamycin and allowed to grow overnight at 30°C and shaking at 200 rpm. 1 mL of the culture was pelleted and plasmid DNA was extracted for sequencing. 500 µL of the grown culture was then used to back dilute into 10 mL of fresh LB supplemented with kanamycin. This back dilution happened 3 times resulting in 4 total time points.

For plate-based selection, plates were incubated overnight at 30 °C. Colonies were then harvested and resuspended in LB. The resuspension was diluted to an OD of 3 and pelleted. The plasmid DNA from this pellet was extracted and used for sequencing. With the same resuspension, 50 mL cultures were started at an OD of 0.5. These cultures were incubated overnight at 30 °C and shaking at 200 rpm and pellets were collected the next day for DNA extraction and sequencing.

### 2.5 Analyzing the plasmid fitness within each bacterial strain

For each sample PCR amplification of the DNA barcodes, and BarSeq, was performed as described previously [21,35]. A list of the count of barcodes per sample was generated. Each file, containing barcode sequencing data for one strain’s time point, was filtered to remove all barcode counts that were below 6. After filtering, an average of 98.64% (± 0.36%) of the barcode counts remained. The filtered barcodes were then compared to the barcode-part association generated and the resulting data was used to generate all origin-related data, as well as the analysis of each recipient bacterium’s preferred part 2s and part 3s. The barcodes for each time point were also investigated for barcode diversity. This was done by determining the number of barcodes representing each origin present at each time point.

### 2.6 Colony quantitative PCR

SsoAdvanced Universal SYBR Green Supermix was ordered from BioRad (BioRad Inc, Hercules, CA). All qPCR primers were selected by using Primer3Plus (https://www.primer3plus.com/index.html) using the qPCR settings. Primers for genomic DNA were ordered for gene IAI47_12350 (a glycosyl transferase/lysophospholipid acyltransferase), a gene found only in the *Pantoea* sp. MT58 genome and not on the introduced plasmid. This primer pair amplifies a region of 528 bp. Primers for plasmid amplification were ordered to amplify a region within the novel origin 6911. This primer pair amplifies a region of 598 bp. Melt curves utilizing all primer pairs show only one amplicon. Plates were run on a CFX96™ Real-Time System using Bio-Rad CFX Manager 3.1 software. A fresh single colony of *Pantoea* sp. MT58 with pAC70, *Pantoea* sp. MT58 with pAC75, and *Pantoea* sp. MT58 with pAC77 were picked into 40 µL of Ambion™ nuclease-free water, separately, and resuspended by pipetting up and down. 20 µL of the resuspended colony was then diluted into 20 µL of water. This was repeated for a total of 3 dilutions (undiluted, 1:2, 1:4, 1:8). 2 µL of each dilution was used for each qPCR reaction, which was performed in triplicate. All colony qPCR data and primers used for all colony qPCR experiments can be found in Data S4.

The thermocycler protocol and qPCR mastermix were executed as described in [36].

Copy numbers (CN) were calculated using the following formula [36]:

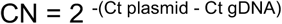

Copy number was then corrected by primer efficiency (E) by the following formula:

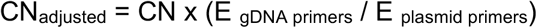

Primer efficiency was calculated using PCR Miner (http://miner.ewindup.cn/miner/), which calculates the primer efficiency of each well by performing a linear regression of the fluorescence values during the exponential phase of amplification.

The final equation used to determine the plasmid copy numbers was the following [37]:

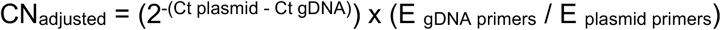

### 2.7 GFP quantification in *Pantoea* sp. MT58

The original magic pool plasmid design utilized a promoter that was non-functional in driving expression of GFP in *Pantoea* sp. MT58. Plasmid pAC80 was constructed combining novel origin 6911, part 2.4 and part 3.8, as well as an inducible promoter, P_BAD_, to drive expression of the GFP. Plasmid pAC80 for GFP expression was conjugated into *Pantoea* sp. MT58 using standard protocols as described above. The empty vector, pAC87, was constructed by removing the GFP from pAC80.

Both the GFP-containing plasmid and the empty vector were conjugated in *Pantoea* sp. MT58 and successful conjugations were observed through colonies resistant to kanamycin. Three colonies of *Pantoea* sp. MT58 containing pAC80 and three colonies of *Pantoea* sp. MT58 containing the empty vector were each inoculated into a 24 deep well plate containing 3 mL of LB with kanamycin and 0.3% (w/v) arabinose and grown overnight at 30 °C while shaking at 200 rpm. After growing overnight, each culture was split into three technical replicates and back diluted (150 µL into 3 mL) in 3 mL of LB with kanamycin and 0.3% (w/v) arabinose for another overnight growth.

1 µL was sampled from the dense overnight culture and added to 99 µL of phosphate buffered saline (PBS) in a 96 well flat bottom plate. GFP was quantified by using a BD C6 Accuri™ flow cytometer by reading 30,000 events using a medium flow rate (35 µL/min, core size = 16 µm). One wash cycle was performed in between each well. Raw data can be found in Data S5. Representative flow cytometry data can be seen in Figure S5.

### 2.8 Indigoidine production

Indigoidine production plasmid pAC75 and empty vector pAC87 were conjugated into *Pantoea* sp. MT58 using standard protocols as described above. A single colony of each strain was inoculated into 5 mL of LB with kanamycin and grown overnight at 30 °C while shaking at 200 rpm. The following day, 300 µL of the seed culture was back diluted into a 24 deep well plate, in triplicate, containing 3 mL of LB with kanamycin and 0.3% (w/v) arabinose. This plate was incubated in a shaker at 30 °C and shaking at 900 rpm. After 24 hours, 200 µL of each sample was taken from the plate and spun down at 14,000 rpm for 5 minutes. The 24 DWP was then put back into the shaker for another 24 hours in which the final time point would be taken. Extraction occurred by removing the supernatant of spun down cultures, resuspending the pellet in 500 µL of DMSO and shaking at 3000 rpm for 20 minutes. Due to the high absorbance of the sample, 20 µL of the extracted material was diluted into 80 µL DMSO and then put into a clear bottom 96 well plate and absorbance was measured at 612 nm (Data S6).

### 2.9 Isoprenol production

Experiments to determine isoprenol toxicity, catabolism and production were performed using minimal salt (M9) medium composed of 1x M9 salts (6.8 g/L Na_2_HPO_4_, 3 g/L KH_2_PO_4_, 0.5 g/L NaCl), 1% (w/v) glucose, 10 mM (NH_4_)_2_SO_4_, 2 mM MgSO_4_, 0.1 mM CaCl_2_ and trace metal solution (500 µL per 1 L medium, Product No. 1001, Teknova Inc, Hollister, CA) [38,39]. All inoculations were grown at 30 °C and shaking at 200 rpm. Isoprenol production plasmid pAC104 was electroporated into *Pantoea* sp. MT58. Electrocompetent cells were prepared by spinning down 2.25 mL of an overnight culture at stationary phase at 8000 rpm for 3 minutes. Supernatant was removed and the pellet was washed with 750 µL of 10% glycerol three times, centrifuging at 8000 rpm for 3 minutes each wash. Washed pellet was resuspended in 250 µL of 10% glycerol. 50 µL of cells was mixed with 50 ng of DNA and electroporated at 2.5 kV/cm in a 1 cm cuvette. Cells recovered in SOC shaking at 900 rpm for 1 hour. 200 µL of cells were plated on LB supplemented with kanamycin and grown overnight.

To determine isoprenol toxicity and catabolism, three single colonies of *Pantoea* sp. MT58 were inoculated in 5 mL LB in glass culture tubes and allowed to grow overnight at 30 °C and shaking at 200 rpm. Cultures were then back diluted in 5 mL M9 with 1% glucose in glass culture tubes and allowed to grow overnight at 30°C and shaking at 200 rpm. This overnight culture was then used as inoculum for further experiments as follows.

For the isoprenol consumption assay, cultures were then inoculated again in M9 (either with or without 1% glucose) either with, or without, 1 g/L isoprenol in a 24 deep well plate at an OD of 0.01, which were sealed and incubated at 30°C with agitation in a microplate reader for 24 hours.

For the isoprenol toxicity assay, cultures were then inoculated again in M9 with 1% glucose and varying concentrations of isoprenol in a 24 deep well plate at an OD of 0.01, which were sealed and incubated at 30°C with agitation in a microplate reader for 24 hours.

All plates were sealed with a semi-permeable film (Breathe Easy Film, USA Scientific, Ocala, FL) and incubated in a microplate reader (Molecule Devices m2E Plate Reader, San Jose, CA). Readings were taken every 15 minutes with continuous agitation at 30 °C in between readings (Figure S7).

For isoprenol production, three single colonies of *Pantoea* sp. MT58 were inoculated in 5 mL of LB supplemented with kanamycin in glass culture tubes and allowed to grow overnight at 30 °C while shaking at 200 rpm. Cultures were then back diluted in 5 mL M9 supplemented with 1% glucose and kanamycin in glass culture tubes and allowed to grow overnight at 30 °C while shaking at 200 rpm. Cultures were then back diluted in 5 mL of M9 supplemented with 1% glucose, kanamycin and either 0.2% (w/v), 1% (w/v) or no arabinose in glass culture tubes and were allowed to grow overnight at 30°C while shaking at 200 rpm. At 24, 48, and 72 hour time points samples were collected to measure cell density (Figure S6) and isoprenol according to [39]. Samples were mixed with equal volumes of ethyl acetate and vortexed at max speed for 15 minutes. Samples were then centrifuged and 80 µL of the upper organic layer was used for analysis using Gas Chromatography-Flame Ionization Detector (GC-FID). 80 μL of the ethyl acetate layer was transferred to a GC vial containing an insert, and 1 μL was analyzed using an Agilent Technologies GC 8890 system (USA) with a FID and a DB-WAX capillary column (15 m × 0.25 mm × 0.25 µm, Agilent Technologies, USA) for quantification. Helium served as the carrier gas at a flow rate of 2.2 mL/min, and the injection volume was 1 μL in splitless mode. The injector and detector temperatures were set to 250 °C and 300 °C, respectively. The GC oven followed a programmed temperature gradient: an initial hold at 40 °C, a first ramping to 100 °C at 15 °C/min and then to 230 °C at 30 °C/min followed by a final hold at 230 °C for 1 min. Data collection and analysis were performed using OpenLab software (Agilent Technologies, USA). Analytical grade standards obtained from Sigma-Aldrich (St. Louis, MO, USA) were used to determine isoprenol concentration.

## 3. Results and Discussion

### 3.1 Environmental plasmids used to predict origins of replication

While there are robust resources for plasmid discovery and analysis, prediction of origins of replication has fewer tools available [7,40,41]. One database for the prediction of origins of replication in the chromosomes of bacteria and archaea is DoriC [8]. Thirty two sub-selected circular plasmids from our earlier plasmidome study were assessed using this tool and led to 18 predicted novel plasmid-based origins of replication that could be experimentally tested [17]. These 18 origins of replication were then synthesized as double strand DNA fragments (see Materials & Methods) on a backbone compatible with the magic pool library construction method.

### 3.2 Origins magic pool library constructed to test origin-host pairings

A magic pool library was constructed using 4 different parts, as outlined in Figure 1. The magic pool was constructed using 38 variants of origins of replication (Part 1), 10 variants of kanamycin drug resistance gene promoters [including RBS] (Part 2), and 2 variants of kanamycin drug resistance genes (Part 3), per the original design of the magic pools plasmid scaffold [21]. Variants of parts 2 and 3 were chosen to ensure effective kanamycin selection in the target microbe. The construction of this library was performed using golden gate assembly and was sequenced using long-read sequencing. In the mapping of the barcode-part association of the magic pool it was found that the colE1-associated plasmids were disproportionately abundant, likely due the high copy number of this origin in *E. coli*.

**Figure 1.**
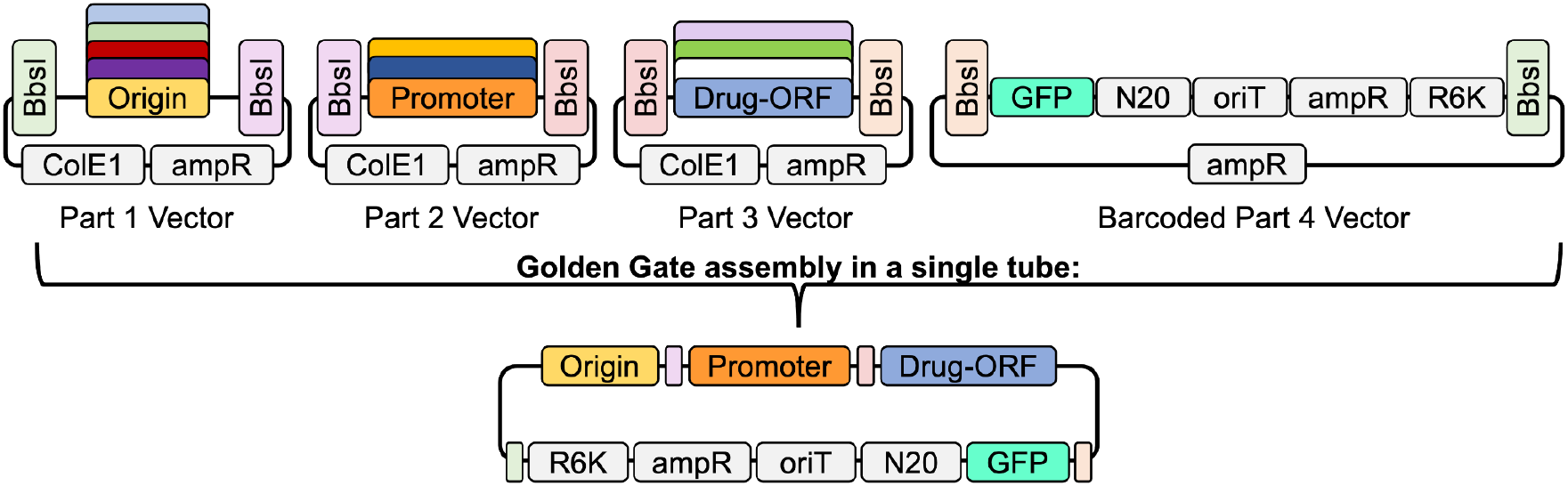
Magic pool construction method. Each colored BsbI box represents a unique sticky end cut site that will complement only the BsbI boxes with the same color to link plasmid parts together sequentially. N20 is representative of the 20 base pair barcodes.

In the library mentioned above, Part 1 is the variable part; the origin sequence and the key component being assessed, represented by one of 18 putative novel origins or one of 20 origins mined from literature (Supplemental Data S2). Parts 2 and 3 came from a previous study [21]. Part 4 was the barcoded backbone for all plasmids constructed in the magic pools and contains a conditional R6K origin of replication for plasmid maintenance in *E. coli*, an origin of transfer, and an antibiotic selection marker (carbenicillin). The sequences of parts 1, 2 and 3 can be found in Supplemental Data S2.

The magic pool backbone (part 4) has a barcode which can establish a plasmid-barcode association. These associations consist of a single barcode and a combination of parts (a single origin, a single promoter, and a single kanamycin drug resistance gene). Barcodes that linked to multiple plasmids or combinations of parts were eliminated from consideration. This resulted in 8,802 total barcoded plasmids that mapped a unique barcode to a part1 origin (some of these barcodes could not be confidently assigned a part 2 or part 3). The number of barcodes associated with each origin in the magic pool can be seen in Figure S1.

### 3.3 Barcode sequencing reveals host-origin pairings

The magic pool library, with variable plasmid origins, was tested for transformation and self-replication capabilities in a set of bacteria including isolates from the ORFRC groundwater. Specifically, *Escherichia coli* BW25113 [42], *Pantoea* sp. MT58 [30] and *Brevundimonas* sp. GW460-12-10-14-LB2 [21] (Data S1: strain and plasmid table) were tested.

To determine if an observable difference could be seen between types of selection methods, two different methods were used. The first method, a plate-based method, collected around 5,000 colonies of post-conjugation recipients on kanamycin-containing agar plates, which contained plasmids from the magic pool. These ∼5,000 colonies were resuspended and used for both plasmid DNA isolation and back diluted into liquid culture tubes. After growing overnight, these liquid cultures were used for DNA extraction and barcode sequencing.

The second selection method utilized an entirely liquid-based method. The bacterial strains used in this study vary in growth rates, which can impact plasmid uptake, replication, retention and partitioning into daughter cells. Due to the difference in growth rate for each strain four consecutive liquid outgrowths to stationary phase followed by dilution were performed to test for the plasmid population stability, setting up a time course (i.e. will one plasmid dominate the others for the entire period or was there a stable mixture of plasmids that can co-exist in the population).

By isolating the plasmid DNA at the end of each outgrowth time point and sequencing the barcode containing PCR product, the magic pool approach allowed us to ascertain which barcodes, and therefore the associated collection of plasmid parts, were enriched. The output of this sequencing data was a list of the barcodes sequenced and the number of reads for each microbe and outgrowth. In all of these experiments, the ability of a recipient cell to grow in the presence of kanamycin depends on multiple factors including whether the origin successfully allows plasmid replication, and a combination of part 2 and part 3 to drive sufficient expression of the kanamycin resistance gene.

Barcode sequencing data was analyzed by comparing the long read sequencing data of the original magic pool, which determined the barcode-origin pairings, to the barcode counts from each dilution. The outcome of this analysis resulted in the counts of each barcode. Barcodes that were enriched by replicating in the host were sequenced more frequently than those barcodes associated with a plasmid unable to replicate in its host. The original full plasmid sequencing which mapped the part combinations to the barcode could then be used to determine the origins that permitted self-replication in a given host microbe (Figure 2).

**Figure 2.**
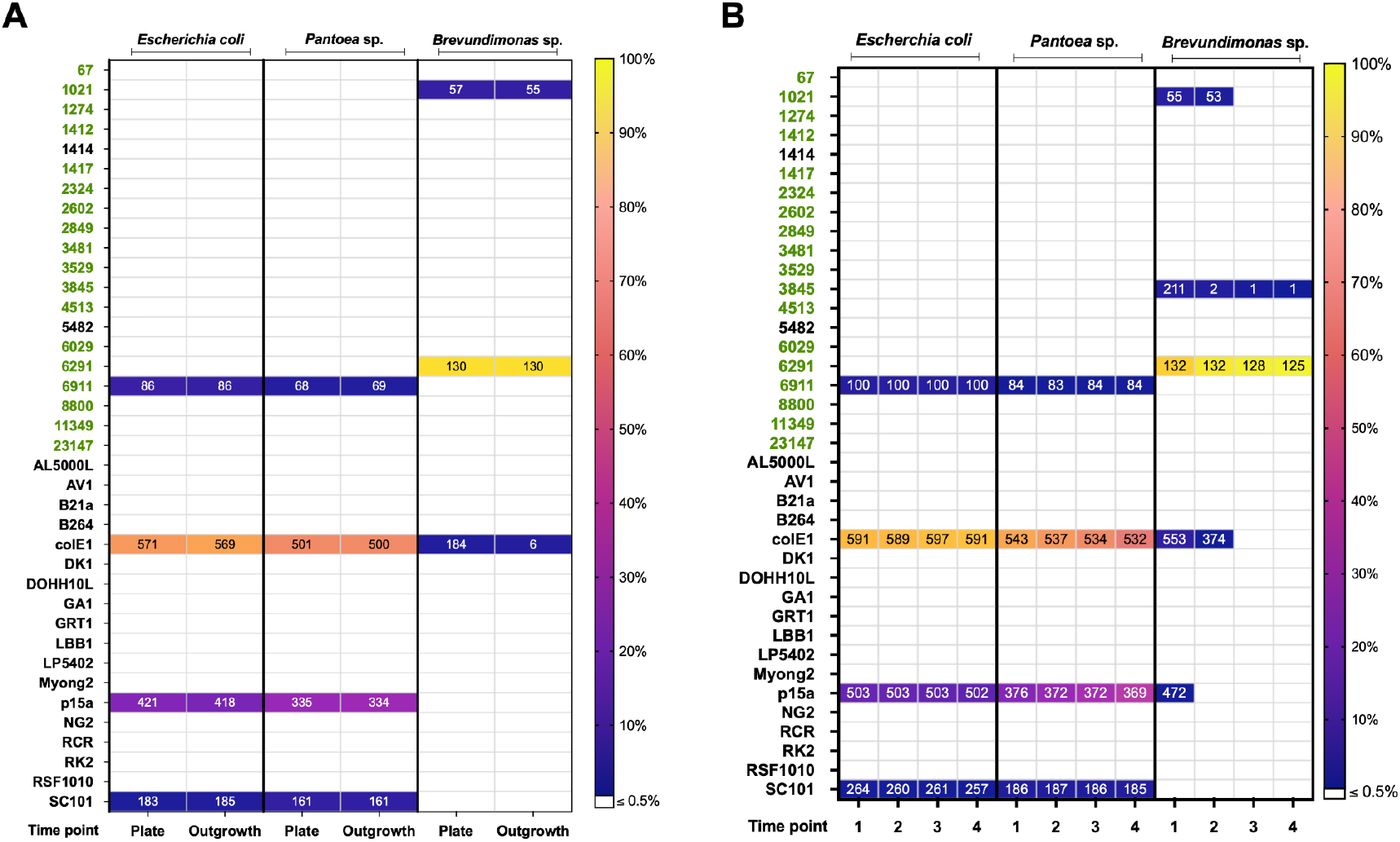
Magic pool BarSeq results. The y-axis denotes the names of the origins of replication, with predicted novel origins in green, and origins found in literature in black. Colors in each time point square represent the fraction of barcode counts associated with that origin versus the total fraction of barcodes in that sample (see legend). The number present in each box represents the total number of unique barcodes detected for that particular origin of replication at that time point. A) Plate-based time points for the magic pool conjugations with the 4 hosts. “Plate” is indicative of cells harvested directly from an agar plate, while “Outgrowth” is indicative of an overnight of growth in liquid media after scraping colonies off of an agar plate. B) Liquid-based time points for magic pool conjugations with the 3 hosts. Time points 1, 2, 3 and 4 represent 24, 48, 72, and 96 hours respectively.

An analysis of the amount of barcodes representing each origin was performed for each time point. We considered an origin to replicate in a given host bacterium if we detected high barcode read abundance, and that the majority of unique barcodes associated with that origin part 1 had high read counts. Similarly, we analyzed the data for high-confidence part 2 and part 3 sequences that worked effectively for driving kanamycin resistance in each strain (see Figures S2-S4 for part 2 and 3 variants breakdown in all strains).

The plate-based method found that 4 origins worked effectively in *E. coli* BW25113 and *Pantoea* sp. MT58 (Figure 2A). Three of these were established origins, p15a, colE1 and SC101, all of which were previously shown to enable plasmid replication in *E. coli* [43–45]. As a new discovery from this study, we found that the novel origin 6911 enabled plasmid replication in both *E. coli* BW25113 and *Pantoea* sp. MT58. In *Brevundimonas* sp. GW460-12-10-14-LB2 it was found that novel origins 1021 and 6291 were capable of conferring plasmid replication. We detected barcode counts from colE1-containing plasmids in this strain, but we do not consider this a functioning origin by our analysis, because only a fraction of the colE1-associated barcodes were detected and barcode counts associated with these origins dropped significantly during the overnight liquid outgrowth.

In the liquid-based magic pool it was found that the results for *E. coli* BW25113 and *Pantoea* sp. MT58 were very similar to the results from the plate-based method, with the same four origins conferring replication, and significant barcode counts observed for all 4 origins even after the last timepoint (Figure 2B). In *Brevundimonas* sp. GW460-12-10-14-LB2, strains with 6291-containing plasmids quickly dominate the entire population, strongly implying that this origin gives the strain a very strong competitive advantage in kanamycin-containing media. Thus, any potential signal from the 1021 origin (which was functional based on the plate data) was drowned out. The barcode counts from plasmids with origins 3845, colE1, and p15a are likely all due to carryover of the *E. coli* donor strain. In support of this, these are all high-abundance parts in the magic pool (Figure S1, S4).

### 3.4 Presence of two origins of replication changes the plasmid copy number in *Pantoea* sp. MT58

Many magic pool generated plasmids containing novel origin 6911 showed evidence of driving plasmid replication in *Pantoea* sp. MT58 and *E. coli* BW25113, based on the BarSeq results (Figure 2). We further investigated this origin of replication and its copy number in the environmentally isolated strain *Pantoea* sp. MT58.

The backbone of the magic pool contains an R6K origin for cloning and for maintaining all of the plasmids in *E. coli* pir+ strains. The existence of this origin of replication in the backbone of a magic pool used to test origins of replication requires the decoupling of the R6K from the origin being tested in the host-origin pair. To decouple the R6K present in the magic pool backbone and the novel origin 6911, and to test the use of this origin, four plasmids were constructed (Figure 3A). These four plasmids were constructed to contain the parts from the magic pool that were necessary (novel 6911 origin and/or conditional R6K origin, kanamycin promoter and resistance gene, and oriT) but also contained a small two-gene pathway to more accurately represent a real world use and application of this plasmid.

**Figure 3.**
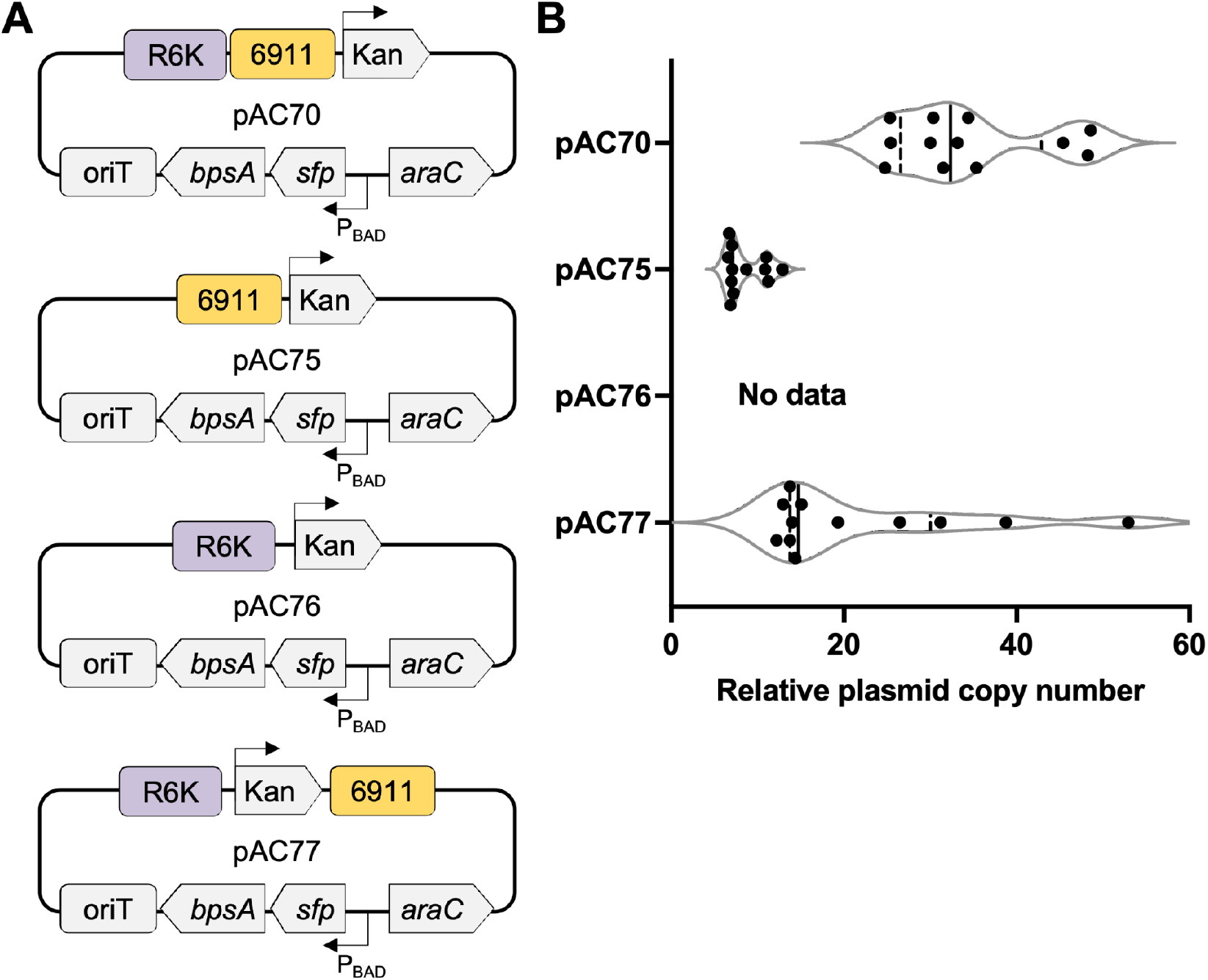
Validation of novel origin 6911 in *Pantoea* sp. MT58. A) Plasmid maps that correspond to the plasmids being tested in (B). B) Violin plot of plasmid copy number. A solid vertical line represents the data median. Vertical dashed lines represent quartiles.

The first plasmid was identical to the plasmid found in the library in regards to the conditional R6K origin and the novel origin 6911 (Figure 3A, plasmid pAC70). This plasmid contains the R6K origin proximal to the novel origin 6911. The second plasmid constructed contains only the origin 6911 as a replication mechanism (Figure 3A, plasmid pAC75). The third plasmid contains only the R6K origin as a replication mechanism (Figure 3A, plasmid pAC76). The final plasmid was constructed with both origins, R6K and 6911, but has placed these origins distal from each other, to see if R6K affects origin 6911 even when not proximal (Figure 3A, plasmid pAC77).

All four plasmids were individually conjugated into *Pantoea* sp. MT58 and successful conjugations were verified by the appearance of kanamycin resistant colonies whereas the control sample did not exhibit spontaneous kanamycin resistance. All plasmids, except for pAC76, were successfully conjugated into *Pantoea* sp. MT58. This indicates that R6K alone was not sufficient for plasmid replication but novel origin 6911 alone was sufficient for plasmid replication in *Pantoea* sp. MT58.

Transformants were then examined using colony-based quantitative PCR (qPCR) to obtain plasmid copy number estimates. qPCR was used to compare the genomic DNA amplicon relative to the plasmid amplicon to provide the number of plasmid copies per genome copy, a relative plasmid copy number per cell, for each plasmid. Rounded to the nearest whole number, pAC70 had a relative copy number of 34 (± 9), pAC75 had a relative copy number of 9 (± 2) and pAC77 had a relative copy number of 22 (± 13) (Figure 3B, Data S4). The method was also tested for a plasmid containing colE1, reported to have 25-30 copies per cell [45]. Colony-qPCR determined that, in *E. coli* 10-β, the plasmid had a copy number of 38 (± 11) (Figure S8). While this experiment queried only origin 6911, the data suggests that having an origin of replication on a plasmid that also contains an R6K origin of replication may affect the plasmid copy number and the plasmid’s replicative capacity in a host.

### 3.5 Expressing a GFP reporter using the novel origin 6911 in *Pantoea* sp. MT58

The backbone of each plasmid in the magic pool encodes a green fluorescent protein (GFP), a commonly used reporter protein in synthetic biology workflows in multiple microbial systems [46,47]. In our study, GFP can be used to test functional origins of replication as well as expression of a heterologous protein in a new host. The original plasmid for *Pantoea* sp. MT58 did not show any GFP expression. However, introduction of an inducible promoter, P_BAD_, (Figure 4A), led to expression of GFP (Figure 4A). Representative flow cytometry samples can be seen in Figure S5.

**Figure 4.**
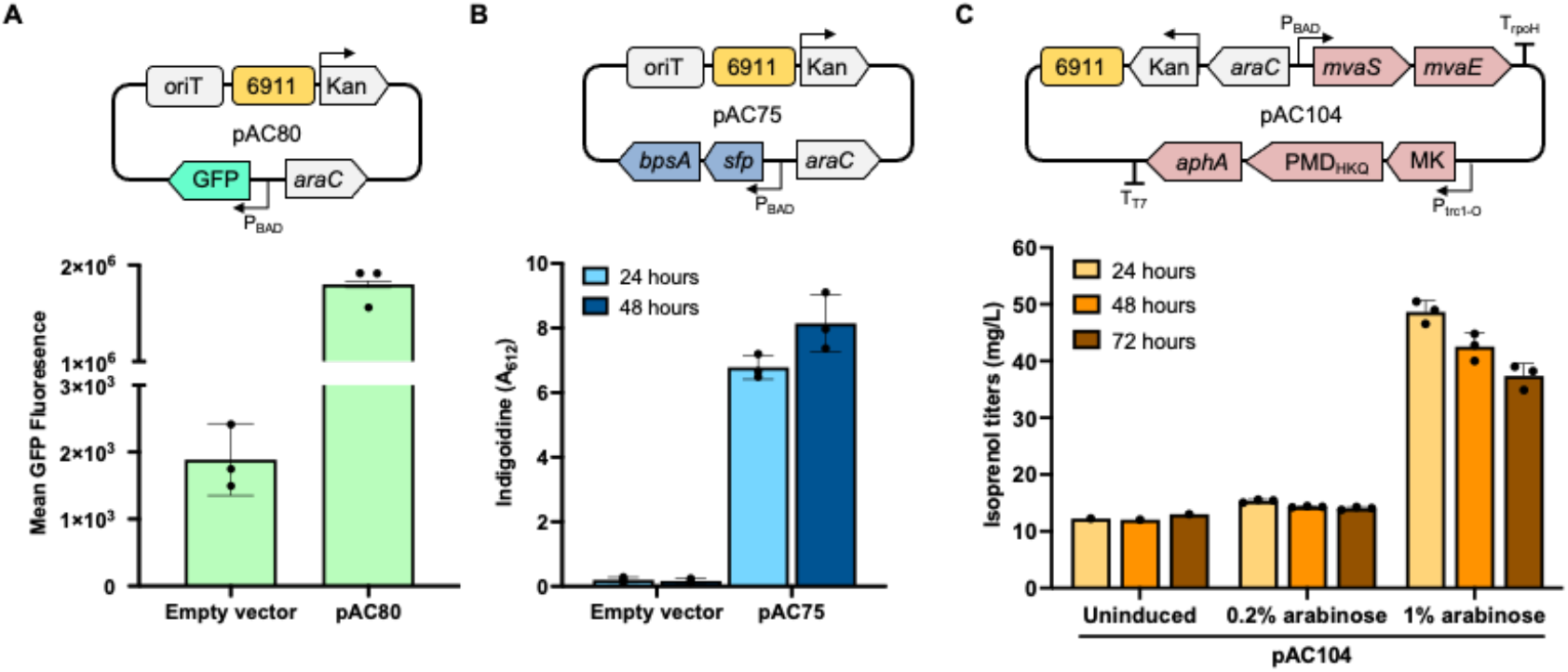
Applications of novel origin 6911 in *Pantoea* sp. MT58. A) *Pantoea* sp. MT58 GFP expression (bottom panel) using plasmid containing novel origin 6911 (top panel). B) Indigoidine absorbance measurements from expression in *Pantoea* sp. MT58 (bottom panel) using a plasmid containing novel origin 6911 (top panel). C) Isoprenol production levels at different inducer levels (bottom panel) using a 6911-containing plasmid (top panel).

### 3.6 Demonstrating the utility of the novel origin 6911 for plasmid-based heterologous pathway expression in *Pantoea* sp. MT58

The successful expression of a reporter gene, GFP, in *Pantoea sp*. MT58 using the novel origin 6911 containing plasmid pAC80 motivated us to further investigate the potential to express pathways for non-native and secondary metabolites. To this end, we transformed *Pantoea* sp. MT58 with arabinose inducible (P_BAD_) multi-gene pathway plasmids, pAC75 and pAC104 in parallel experiments and produced a non-native non-ribisomal peptide product indigoidine, and the hemiterpene compound isoprenol, respectively (Figure S9).

In the engineered *Pantoea* strain heterologously expressing *sfp*_*B. subtilis*_::*bpsA*_*S. lavendulae*_ on the plasmid pAC75, upon induction by 0.3% (w/v) arabinose and extraction of harvested cells with DMSO, we observed an average OD_612_ of 4.4 at 24 hours and 5.0 at 48 hours (Figure 4B, Data S6). The OD_612_ values are indirect measures of indigoidine titers and confirmed expression of the two-step indigoidine pathway containing the phosphopantetheinyl transferase (sfp) and the non-ribosomal peptide synthetase (NRPS, BpsA). Successful production of indigodine represents an ability to express a non-native natural product in *Pantoea sp*. MT58 leveraging the plasmid-based expression system enabled by the novel origin 6911. Indigoidine is also a valuable bio-based synthetic indigo alternative that has been previously reported in other industrially relevant hosts [48–52].

We also validated the use of the novel origin 6911 based plasmid to express products from more complex multi-gene pathways. For this we used the heterologous isoprenoid pathway for the hemiterpene short chain alcohol isoprenol. A previously reported plasmid pIY670 [39] was converted into pAC104 by swapping the origin RK2 (optimized for expression in *P. putida*) with the novel origin 6911. Terpenes span a wide array of compounds from those useful in altering the metabolic profile of a microbe to those that serve as bioproduction targets. In this case, isoprenol is also a useful commodity chemical and a well established biofuel precursor [39,53]. Short chain alcohols also serve important roles in fitness and interactions in microbial communities [54–57] and an ability to express them provides additional tools to alter and examine microbial systems in context of their community. In our system, we observed a maximum isoprenol production of 48.67 mg/L in 24 hours (Figure 4C, Data S7) in glucose minimal medium. A drop in isoprenol titers (to 37.38 mg/L) is observed by 72 h and consistent with published reports [39]. *Pantoea* sp. MT58 cannot utilize isoprenol as the sole carbon source (Figure S7A), however, this strain encodes many alcohol dehydrogenases [58], that could degrade isoprenol. Hence, the decrease in titers at 72 h in the engineered *Pantoea* sp. MT58 could be attributed to either the inherent volatility of isoprenol and/or non-catabolic degradation.

Between the two non-native systems, inducer requirement was markedly higher to achieve metabolite production in the isoprenoid system (1% w/v arabinose) vs. that from the NRPS system for indigoidine (0.3% w/v arabinose) (Figure 4B and C). While indigoidine is produced from the amino acid precursor–glutamate, isoprenol is derived from acetyl-CoA which is a key central metabolite at the intersection of carbohydrate, lipid, and protein metabolism with higher competing pathway demand. Additionally, the heterologous isoprenol pathway is also cofactor-dependent [39] compared to the indigoidine pathway which could impose a metabolic burden. The accumulation of pathway intermediates may impose additional toxicity even though *Pantoea* sp. MT58 is tolerant to up to 7 g/L of the final product–isoprenol (Figure S7B) [59–61]. Furthermore, *Pantoea* sp. MT58 consumes arabinose as a sole carbon source (Figure S10). These results establish the novel origin 6911 as a promising genetic tool that improved the genetic tractability of *Pantoea* sp. MT58, and are an important first step in strain domestication to investigate the ecological role of this bacterium and to develop industrially relevant hosts. However additional optimizations are needed to use these systems to either modulate the levels of these metabolites to examine their role in community interactions, or to facilitate the development of chassis strains.

## 4. Conclusions

Discovering host-origin pairings is essential to moving from genomics-to genetics-based approaches. Our workflow identified a novel origin pairing in the non-model *Pantoea* sp. MT58, which was confirmed through qPCR experiments by decoupling of the conditional R6K origin and the novel origin 6911. The low copy number origin 6911 was then used to express three molecules, each with increasing complexity, in *Pantoea* sp. MT58. While the magic pools library was used to screen a small subset of hosts, both model and non-model, it can be expanded to include additional bacterial hosts for further discovery. The products described in this study are highly relevant to key metabolic pathways, and production of chemicals and fuels. Overall the development of genetic tools for this microbial host has value in its use as a biotechnology platform as well as in future studies for understanding its role in the environment.

## Supporting information

Codik et al 2025 SI

## Acknowledgements

The authors thank the Mukhopadhyay group for their constructive feedback regarding the manuscript. Dr. Shweta Priya (LBNL) provided valuable guidance in conducting the qPCR experiments. We thank the authors of DoriC for initial help with assessing putative origins of replication. We also thank the JBEI (jbei.org) strain collection for two of the strains used in this manuscript.

This work was part of the Ecosystems and Networks Integrated with Genes and Molecular Assemblies (ENIGMA) (http://enigma.lbl.gov) project, a Science Focus Area Program at Lawrence Berkeley National Laboratory (LBNL), and is supported by the U.S. Department of Energy, Office of Science, Biological and Environmental Research program, under contract number DE-AC02-05CH11231 between LBNL and the U.S. Department of Energy. The U.S. Government retains and the publisher, by accepting the article for publication, acknowledges that the U.S. Government retains a non-exclusive, paid-up, irrevocable, worldwide license to publish or reproduce the published form of this manuscript or allow others to do so, for U.S. Government purposes.

## Author Contributions

AC, AK and AM developed the study. AK, HL and AMD designed the origins magic pool library. AC and TO built the library. AC and BLW tested the library. AC performed qPCR, GFP and indigoidine experiments. AR developed electroporation methods for *Pantoea* sp. MT58. AC and AS performed isoprenol experiments, with AS performing GC-FID data analysis. AC performed all other data analysis. TE, AMD and AM provided supervisory roles and feedback. APA, AMD, AM acquired the funds for the project. AC drafted the initial manuscript. All authors have read, provided feedback, and approved the manuscript for publication.

