## Supplementary material for "Novel origin of replication for environmentally isolated *Pantoea* strain enables expression of heterologous proteins, pathways and products": Codik et al 2025 SI

### Abstract

Plasmids isolated or characterized from environmental samples serve as a resource that can be used to develop genetic tools for characterizing recently isolated or less-studied microbes. In this report, we leveraged sequences from a previously characterized groundwater plasmidome and developed a screen to identify novel plasmid origins. Putative origin sequences were used to construct a barcoded plasmid library, which contained both known and newly predicted origins. This library was tested against a panel of representative bacterial strains and led to the identification of 3 novel origins that putatively replicate in gram-negative bacteria not previously associated with these origin sequences. We empirically validated one of the newly identified origins, 6911, to be functional in both the model bacterial strain, *Escherichia coli* BW25113, as well as in *Pantoea* sp. MT58, a fast growing and metal tolerant, environmentally important bacterium from the widespread *Pantoea* genus. We confirmed that a plasmid bearing origin 6911 as the sole origin could replicate and had a copy number of 9 (± 2) in *Pantoea* sp. MT58. We successfully used a plasmid based on the new origin to express the reporter protein GFP, and two non-native metabolite pathways for the natural product, indigoidine and the terpenoid compound, isoprenol. By pairing functional novel origins of replication to non-model organisms this pipeline can expand the tool kit for genetic manipulations of both model and less-studied bacteria.

**Supplemental information table of contents:**

- Strain and plasmid table - Data_S1.xlsx
- Parts and their sequences - Data_S2.xlsx
- Barcode-part combinations - Data_S3.xlsx
- qPCR raw data/primers used - Data_S4.xlsx
- GFP expression - Data_S5.xlsx
- Indigoidine absorbance data - Data_S6.xlsx
- Isoprenol expression data - Data_S7.xlsx

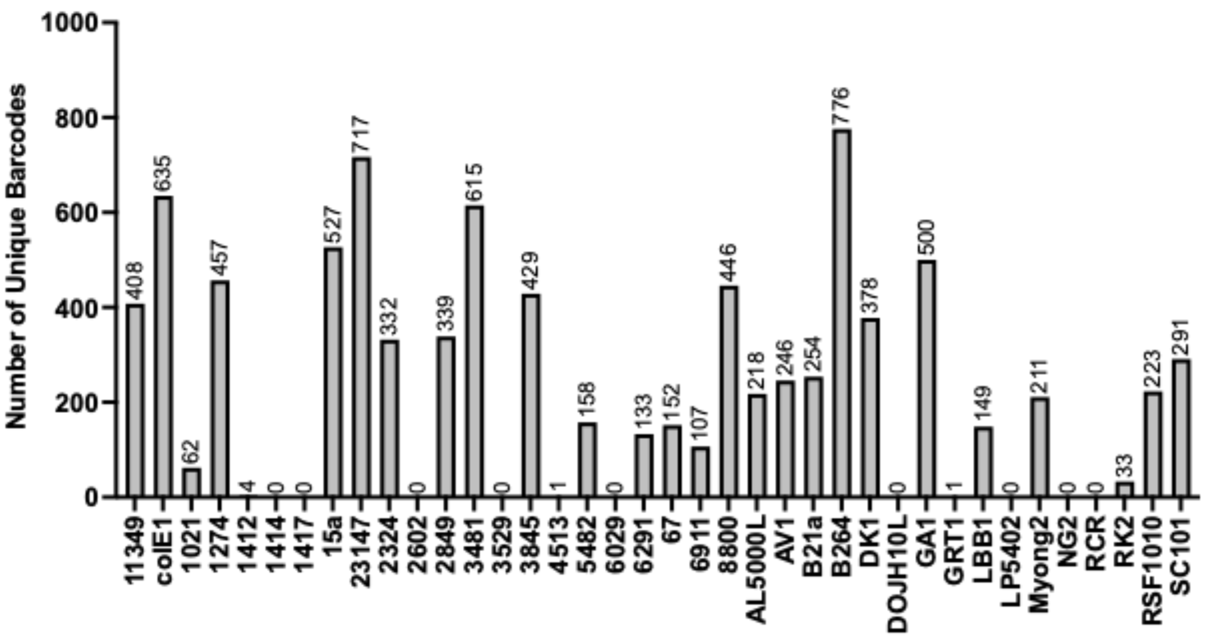


Figure S1. Number of unique barcodes per origin present in the magic pool.

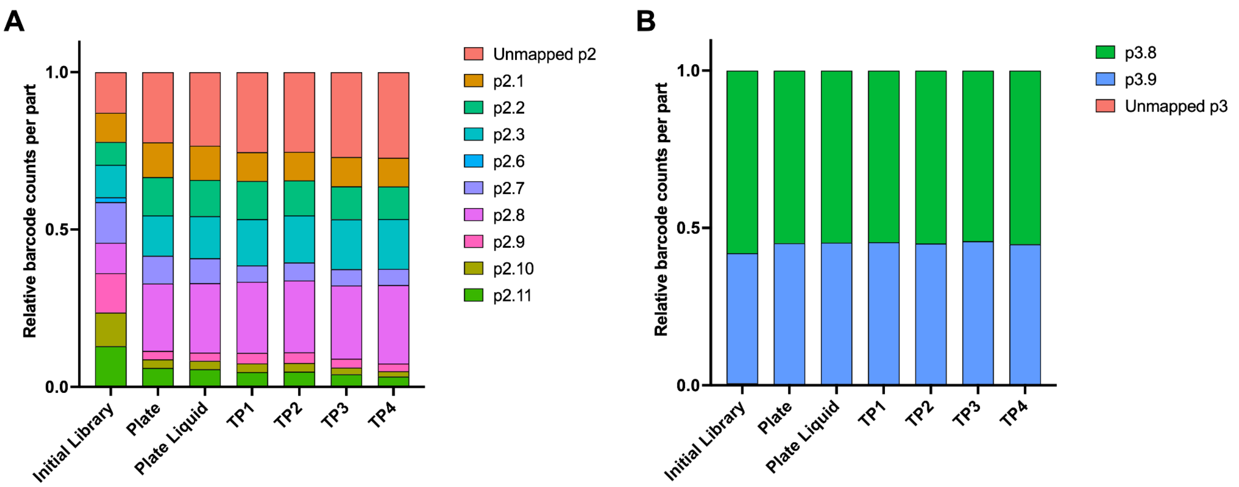


Figure S2. Breakdown of part 2 and 3 variants in *E. coli* BW25113.

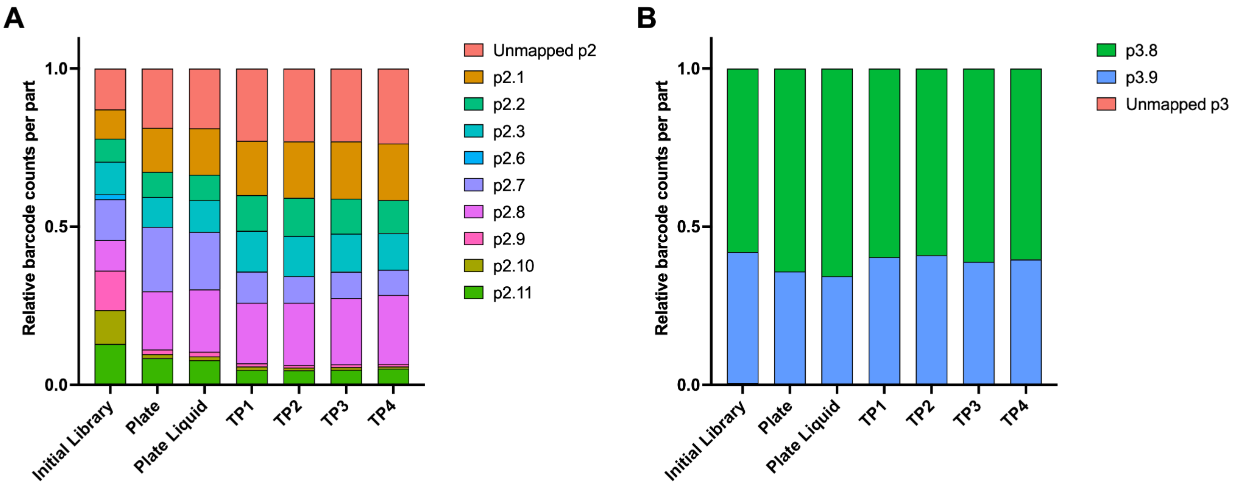


Figure S3. Breakdown of part 2 and 3 variants in *Pantoea* sp. MT58.

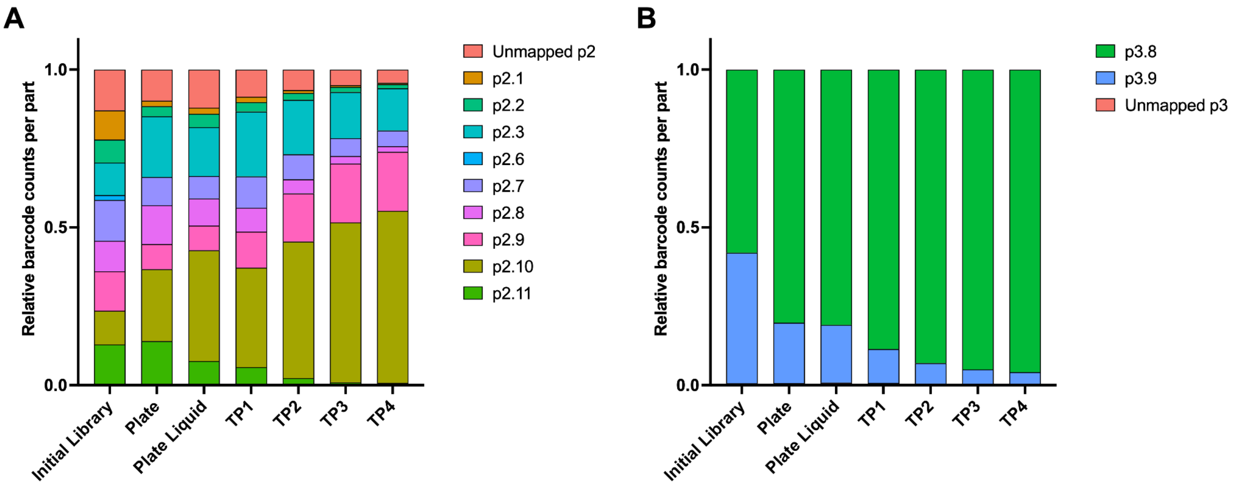

Figure S4. Breakdown of part 2 and 3 variants in *Brevundimonas* sp. GW460-12-10-14-LB2.

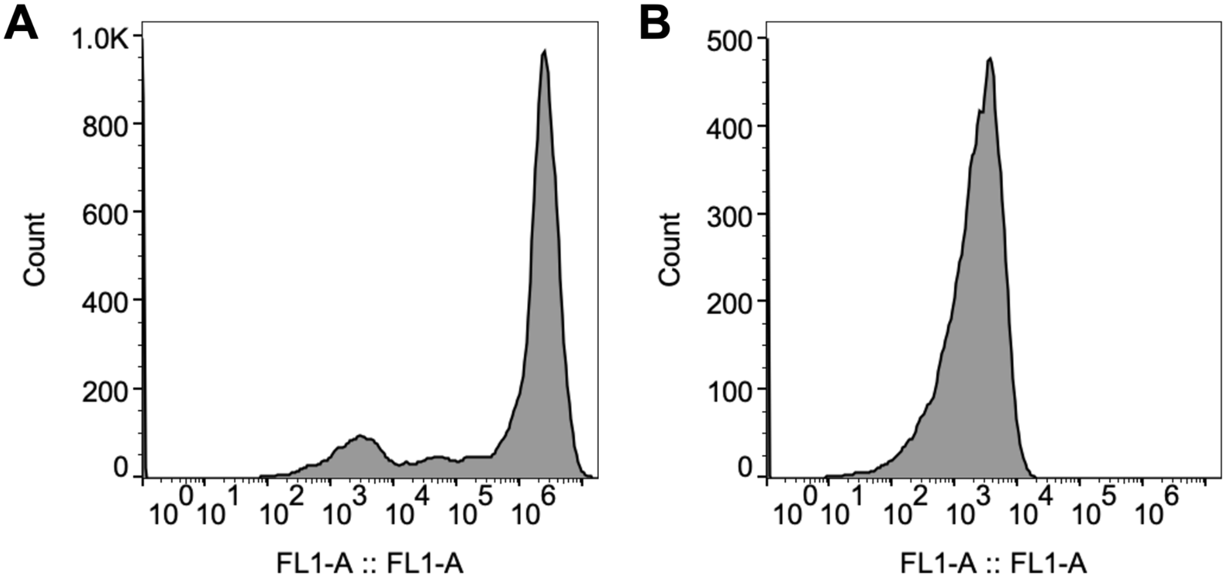


Figure S5. Flow cytometry GFP expression data. A) Representative fluorescence data from GFP expressing *Pantoea* sp. MT58 cells. B) Representative fluorescence data from *Pantoea* sp. MT58 cells without the GFP-expressing plasmid.

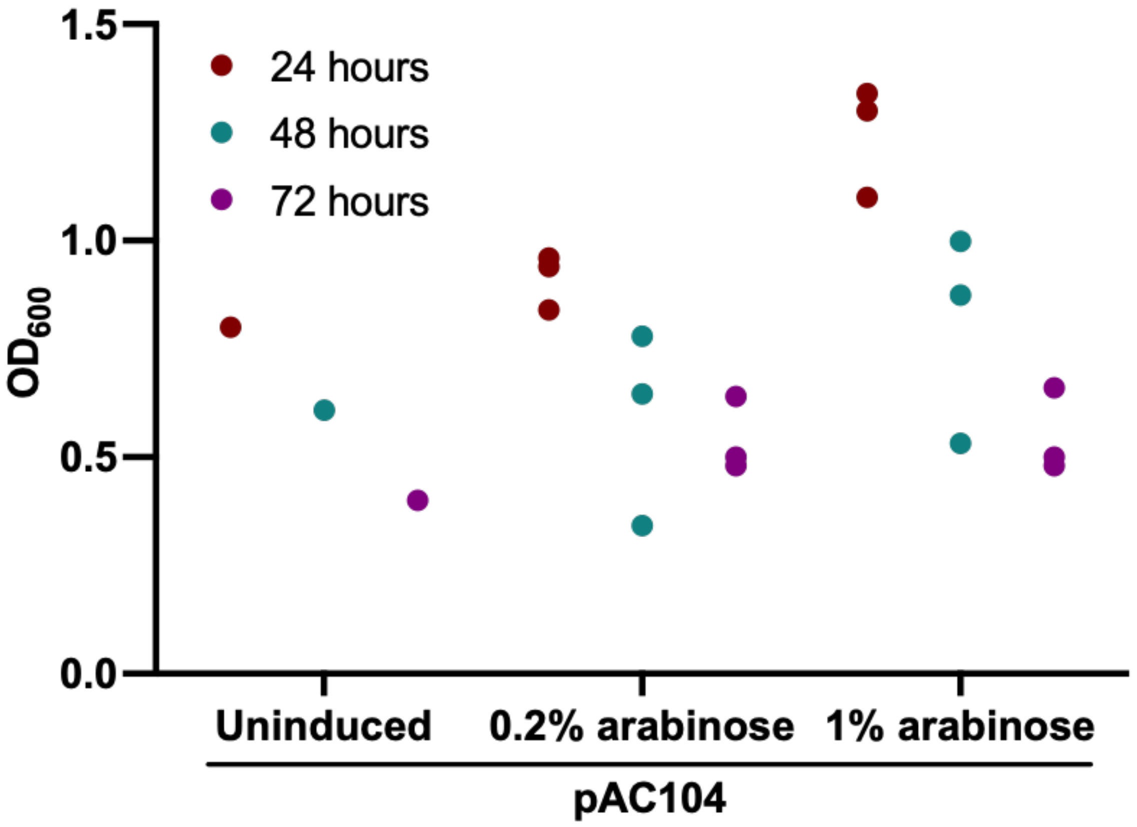


Figure S6. Optical density measurements for isoprenol production run.

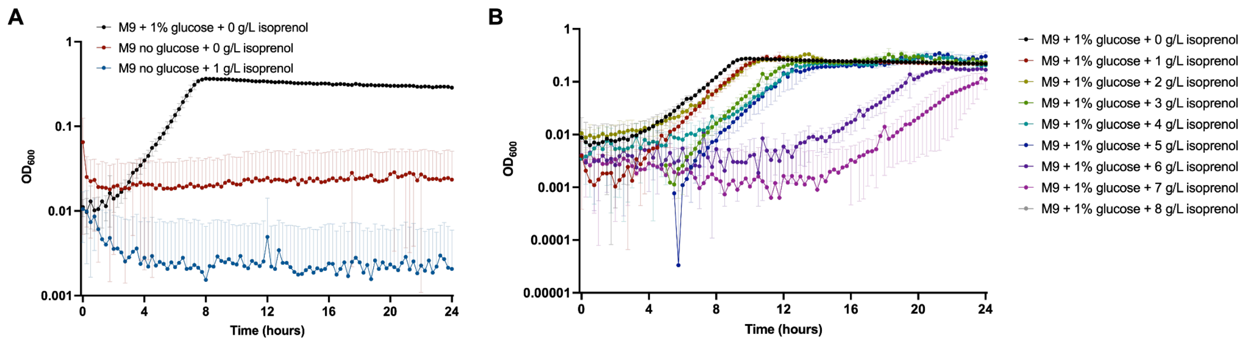


Figure S7. Isoprenol degradation and tolerance profile in *Pantoea* sp. MT58 A) Determination of *Pantoea* sp. MT58's ability to consume isoprenol as a sole carbon source. B) *Pantoea* sp. MT58's growth response to varying concentrations of isoprenol supplemented with M9 1% glucose minimal medium. The data for M9 + 1% glucose + 8 g/L isoprenol, when corrected against the blank, resulted in negative values, and are indicative of a lack of growth.

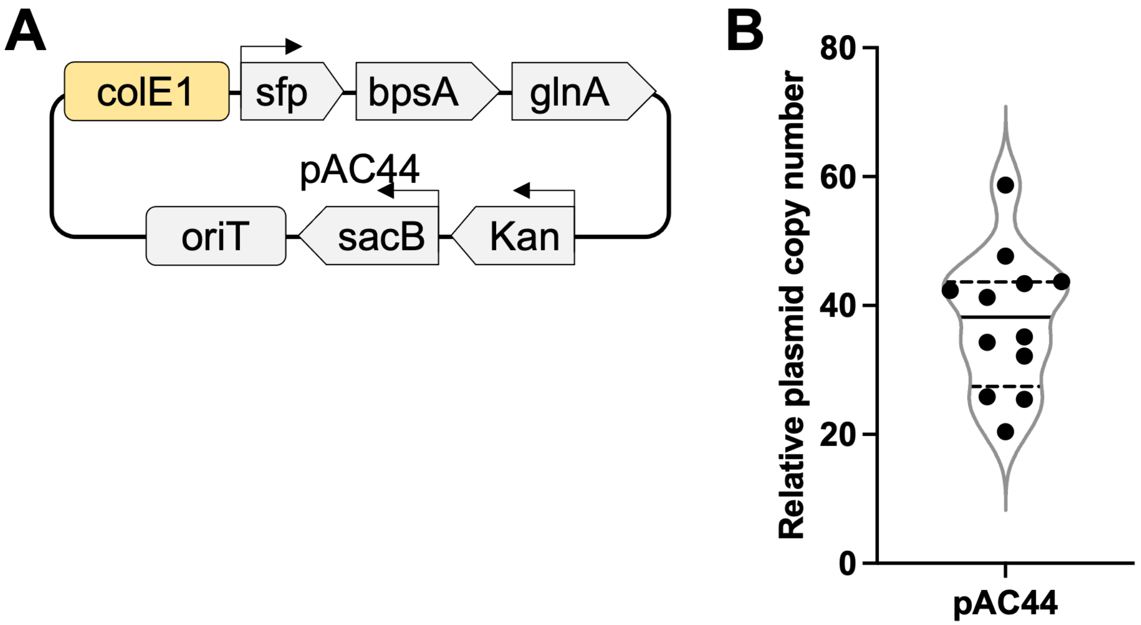


Figure S8. Validation of colony qPCR method through testing known colE1 origin. A) Plasmid map of plasmid used to generate qPCR data in (B). B) Violin plot of plasmid copy number. A solid vertical line represents the data median. Vertical dashed lines represent quartiles.

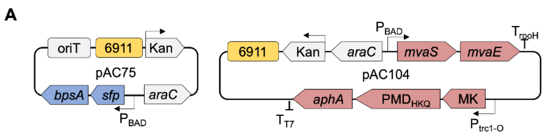


**B**

| Indigoidine pathway (pAC75) | | Isoprenol pathway (pAC104) | |
|---|---|---|---|
| **Gene name** | **Gene Description** | **Gene name** | **Gene Description** |
| *bpsA* | Blue pigment synthetase A | *mvaS* | hydroxymethylglutaryl-CoA synthase |
| *sfp* | PPTase | *mvaE* | acetyl-CoA acetyltransferase |
| | | MK | mevalonate kinase |
| | | $PMD_{HKQ}$ | phosphomevalonate decarboxylase |
| | | aphA | Aminoglycoside 3'-phosphotransferase |

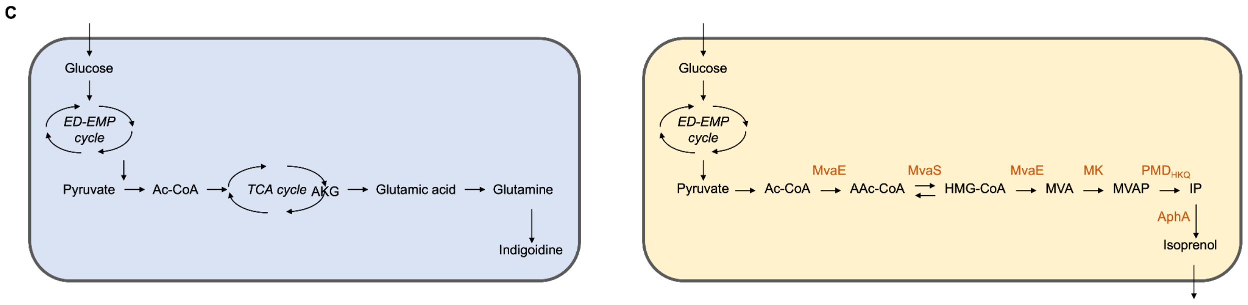


Figure S9. Indigoidine and isoprenol pathway breakdowns. A) Plasmid maps of plasmids used to express indigoidine (left panel) and isoprenol (right panel) in *Pantoea* sp. MT58. B) Gene name and description of genes used in each plasmid for expression of indigoidine and isoprenol. C) Representative cell map and metabolisms to produce indigoidine (left panel) and isoprenol (right panel).

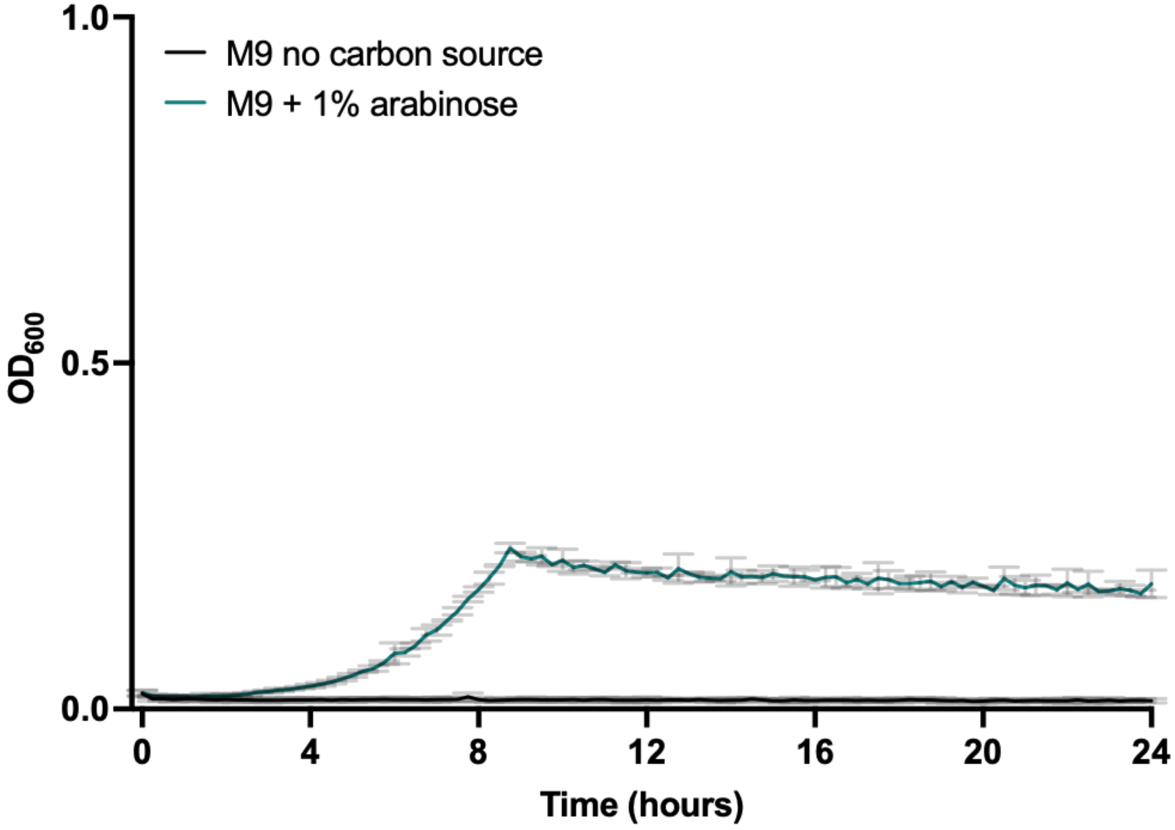


Figure S10. *Pantoea* sp. MT58 growth on arabinose as a sole carbon source. Cells grown in a 96 well plate. N = 3
